# Temporal regulation of human reactive astrocytes reveals their capacity for antigen presentation

**DOI:** 10.1101/2025.06.26.661787

**Authors:** Emily Jane Hill, Caitlin Sojka, Maureen McGuirk Sampson, Hsiao Lin Wang, Alexia Tyler King, Anson David Sing, Emory Brain Organoid Hub, Victor Faundez, Steven A. Sloan

## Abstract

Astrocytes adapt to injury and disease by entering a reactive state defined by transcriptomic, morphological, and functional changes. Using a combination of human cortical organoids (hCOs) and primary fetal brain tissue, we investigated the plasticity of human astrocyte reactivity. We observed robust inflammatory transcriptomic and chromatin signatures following cytokine exposure, which varied with duration. To assess reversibility, we withdrew cytokines after acute or chronic exposure. In both cases, astrocytes returned to a quiescent genomic state within days. Chronic exposure induced MHC class II gene expression, normally restricted to professional antigen-presenting cells. We validated MHCII protein in primary tissue and hCOs and used co-immunoprecipitation and mass spectrometry to identify candidate antigens. Finally, we showed that exogenous peptides from fetal neurons could be presented by astrocytic MHCII.

## Introduction

Astrocytes perform a wide range of neurodevelopmental and neuroprotective functions related to synaptic formation and function (1, 2), neurotransmitter and metabolite recycling (3, 4), and blood-brain barrier structure and maintenance (5, 6). In addition to these roles, astrocytes exhibit remarkable plasticity in response to disease or injury. Early studies categorized reactive astrocyte states into binary groupings (7, 8), but recent work has revealed a much broader spectrum of reactive phenotypes (9). It is now well established that reactive astrocytes display extensive cell state heterogeneity, with responses tailored to the specific nature of the insult or disease context (9). Advances in single cell/nucleus RNA sequencing have enabled in-depth transcriptional profiling of reactive and disease-associated astrocytes, both in post-mortem human tissues and in vivo murine disease models (10–13). Despite these efforts, relatively little is known about the temporal dynamics of astrocyte reactivity—specifically, the extent to which reactive states can be reversed over time. Previous studies in rodents suggested that acutely reactive astrocytes may return to quiescence after a single, acute insult (14,15). However, many triggers of astrocyte reactivity in humans are prolonged and persistent (e.g. low levels of chronic inflammation and/or neurodegenerative settings). This raises several key questions: How might human reactive astrocytes differ after a chronic insult compared to an acute stimulus? Can both acutely- and chronically-activated reactive astrocytes return to a quiescent state after injury? Answering these questions will be critical in fully understanding reactive astrocyte contributions and functions in disease, as well as how they may be targeted or manipulated.

Seminal work from the Barres and Liddelow labs demonstrated that a combination of microglial-secreted inflammatory cytokines, TNF-α, IL-1α, and C1q (collectively referred to as TIC) (8, 16) is sufficient to induce a canonical inflammatory astrocyte state. Transcriptionally, this response is marked by a rapid (within 24 hours) upregulation of inflammatory signaling cascades, such as the JAK/STAT and NF-κB pathways, and the release of inflammatory molecules including IL-6 and CXCL10 (7, 8, 17–19) Notably, this inflammatory signature is conserved across species: canonical TIC-induced genes are also robustly upregulated in 2D human astrocyte cultures following a single exposure (18, 19), and similar transcriptional profiles have been observed in astrocytes from post-mortem human brain tissue across a range of neurological disease states (10, 11, 13, 20, 21). These findings suggest that TIC induces a relatively core inflammatory astrocyte response, which is subsequently colored by specific genetic vulnerabilities and/or extrinsic disease states. However, many inflammatory states are not transient (13, 22–24), and little is known about the temporal dynamics of reactive astrocyte states as they transition from acute to chronic stages of inflammation. Understanding how the reactive state is sustained or reshaped over time is essential for elucidating the full range of astrocyte contributions in disease and identifying potential points for therapeutic intervention.

The development of advanced in vitro cell culture systems, both in 2D and 3D, has allowed for the modeling and study of CNS cell types over time and in the context of disease (25–30). To investigate how astrocytes transition from the initiation of reactivity to a chronically inflamed state, we leveraged the human cortical organoid (hCO) model system (27, 30). hCOs are three-dimensional cell cultures that recapitulate key features of developing cortical cytoarchitecture, are amenable to long-term culture, and can be readily manipulated for experimental perturbation (31–33).

To investigate the temporal dynamics of astrocyte reactivity, we treated hCOs with the TIC cytokine cocktail for durations ranging from 1 day to 3 months. We then assessed both chromatin accessibility (via ATAC-seq) and transcriptional responses (via RNA-seq) specifically in astrocytes. We observed distinct, time-dependent molecular responses to acute versus chronic TIC exposure. Upon TIC withdrawal, we found that astrocytes largely reverted to a baseline transcriptional state, even after prolonged exposure. Notably, chronic exposure also revealed a previously unrecognized astrocyte response: robust induction of major histocompatibility complex class II (MHCII) genes and protein expression. As astrocytes are not canonical anti-gen-presenting cells, this finding was unexpected. We validated MHCII induction using multiple orthogonal approaches at both RNA and protein levels and confirmed surface presentation of MHCII-antigen complexes on astrocytes. To rule out the possibility that this was an artifact of the organoid system—which lacks endogenous myeloid cells—we replicated these findings in ex vivo human fetal brain tissue exposed to TIC, confirming that MHCII induction is a bona fide astrocyte response to chronic inflammatory signaling.

## Results

### Exposure to inflammatory cytokines induces reactive astrocytes in hCOs

The cytokine trio of TNF-α, C1q, and IL-1α (TIC) robustly induces the formation of inflammatory astrocytes in mice (Liddelow et al. 2017), as well as in human astrocyte cultures (18, 19). To examine the temporal dynamics of inflammatory astrocyte responses in a more physiologically relevant human system, we asked whether TIC exposure could elicit a similar inflammatory phenotype in endogenously generated astrocytes within human cortical organoids (hCOs). This required first purifying astrocytes from hCOs following TIC exposure. Because astrogenesis begins around day 90 in hCOs (31, 34) and astrocytes continue to mature with extended culture (33, 60), we performed the following studies using hCOs aged 175 to 250 days, when astrocytes exhibit more mature phenotypes. We purified astrocytes using immunopanning, first negatively depleting neurons (CD24+), and then positively selecting astrocytes (HepaCAM+). We confirmed the purity of both cell populations through bulk RNA-sequencing, which confirmed clear enrichment of neuronal and astrocyte marker genes, respectively (**Fig. 1A**). We next performed bulk RNA-sequencing of astrocytes immunopanned from hCOs exposed to a brief 24-hour pulse of TIC cytokines and observed a robust induction of inflammatory genes, including canonical markers C3, NFKB, and IL-6 (**Fig. 1B**). We also confirmed that TIC exposure in hCOs induced C3 mRNA expression via RNAscope, noting a robust increase in C3 puncta compared to a nearly absent signal in control hCOs (**Fig. 1C**). To test whether these inflammatory signatures were specific to organoids, we treated primary fetal human astrocytes, immunopanned from gestational week (GW) 19-20 cortices, with TIC for 24 hours and found substantial overlap in DEGs, reassuringly mirroring the response in hCO-derived astrocytes (**Supp. Fig. 1A-C**). We next benchmarked the transcriptional concordance of our inflammatory astrocytes in hCOs to existing monoculture studies of hiPSC-derived astrocytes (18, 19) (**Fig. 1D**). While all datasets showed strong concordance in inflammatory gene upregulation, hCO astrocytes consistently expressed higher levels of maturation-associated genes (e.g., AQP4, SLC1A3), likely reflecting differences in astrocyte maturity between 2D and 3D models. Having confirmed robust astrocyte reactivity in TIC-exposed hCO cultures, we next examined the transcriptional heterogeneity of this response using single-cell RNA-seq on control hCOs and those exposed to TIC for either 1 or 7 days (**Fig. 1E**). We observed a largely homogeneous population of inflammatory astrocytes, with the proportion of reactive cells increasing in a time-dependent manner (**Fig. 1F-H**).

**Fig. 1.**
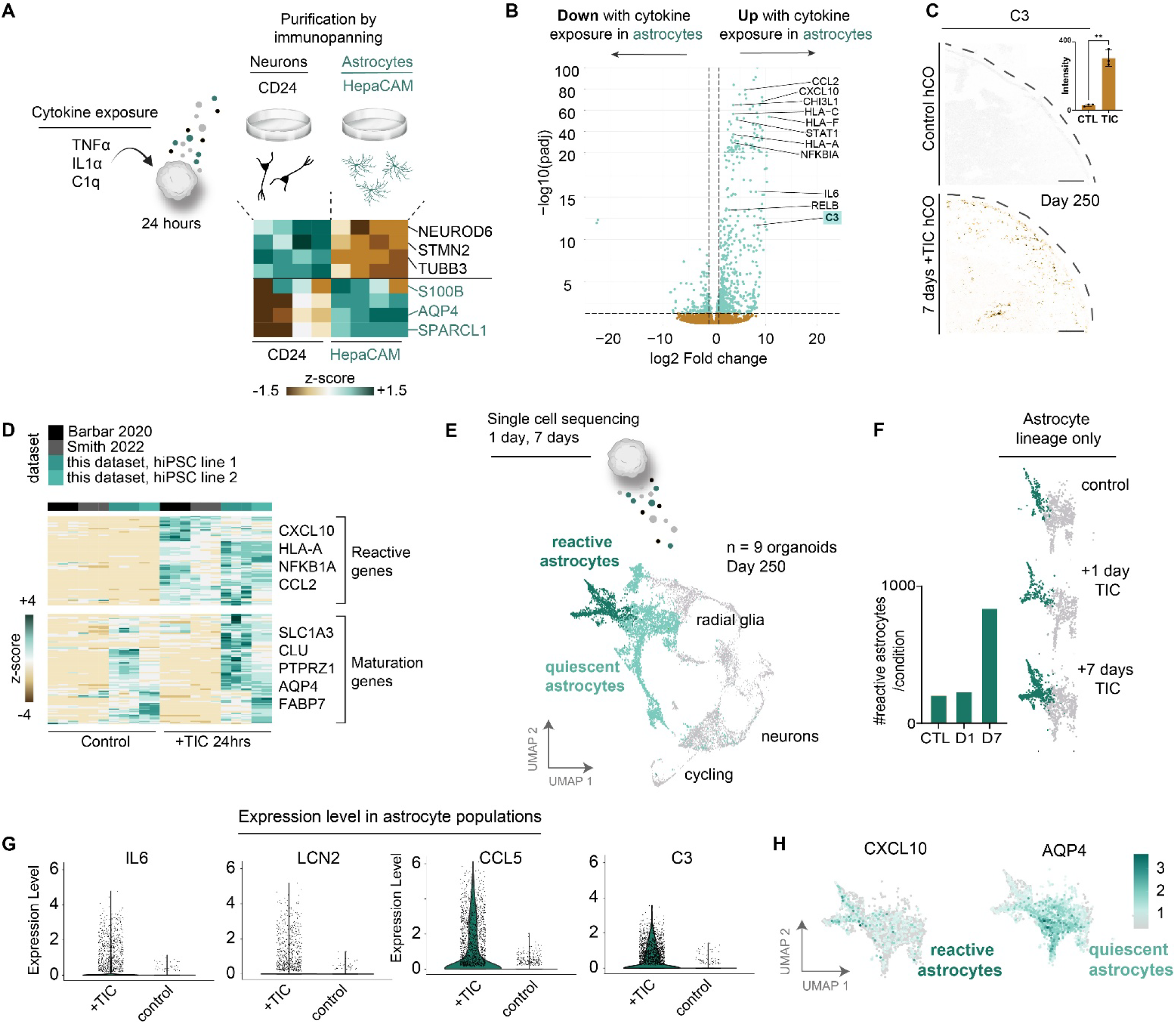
TIC exposure induces robust transcriptional activation in human astrocytes. **(A)** hCOs were exposed to TNF-a, IL-1a, C1q (TIC) for 24hrs. Both neurons and astrocytes were purified from hCOs via immunopanning using antibodies against CD24 and HepaCAM, respectively. Purification was confirmed by bulk RNA-seq via the expression of canonical cell type-specific markers. **(B)** Volcano plot of genes upregulated after 24hrs +TIC in astrocytes. **(C)** RNAscope of C3 mRNA in control (unexposed) or 7 days +TIC hCOs sections (D211, scale= 500 µm). **(D)** Hierarchical clustering of top differentially expressed genes between 24hr +TIC hCOs astrocytes (n = 2 iPSC lines, 8-12 hCOs/line) and 24hr +TIC astrocytes grown in 2D cultures from two previously published datasets (Barbar 2020 and Smith 2022). **(E)** Compiled UMAP of 12,800 cells from unexposed, 24hrs +TIC, and 7 days +TIC hCOs. A population of reactive astrocytes was defined by expression of canonical inflammatory genes. n=3 hCOs/condition. **(F)** UMAP of astrocyte subcluster from E separated into samples from unexposed, 24hrs +TIC, and 7 days +TIC hCOs. **(G)** Violin plots of expression of classical inflammatory reactive astrocyte genes in the astrocyte cluster. **(H)** Feature plots showing cell distribution and expression of CXCL10 (reactive astrocyte marker) or AQP4 (mature astrocyte marker).

### Temporal dynamics of inflammatory reactive astrocytes at genomic scales

Given the time-dependent increase in reactive astrocyte proportions within TIC-exposed hCOs, we next investigated how astrocyte transcriptional and epigenetic signatures change depending on the duration of the inflammatory exposure. To investigate the temporal dynamics of inflammatory human astrocytes, we exposed hCOs to TIC for 1, 7, 31, or 84 days (spanning from acute to chronic exposure) and performed paired ATAC- and RNA-seq on purified astrocytes from both TIC-treated and age-matched control hCOs at each timepoint (**Fig. 2A**). Using weighted gene co-expression network analysis (WGCNA), we identified three major genomic modules shared across both RNA and ATAC datasets. These include (1) a downregulated module comprised of genes and regulatory regions consistently suppressed across all timepoints (88 DEGs / 597 DARs); (2) a time-independent module containing acutely induced genes and accessible regions activated within one day and stably maintained through chronic exposure (1101 DEGs / 9857 DARs); and (3), a time-dependent module defined by genes and regions gradually upregulated or opened over prolonged exposure, many requiring over one month to reach peak expression/accessibility (255 DEGs / 5085 DARs) (**Fig. 2B–C, Supp. Fig. 3, Supp. Data 1-3**).

**Fig. 2.**
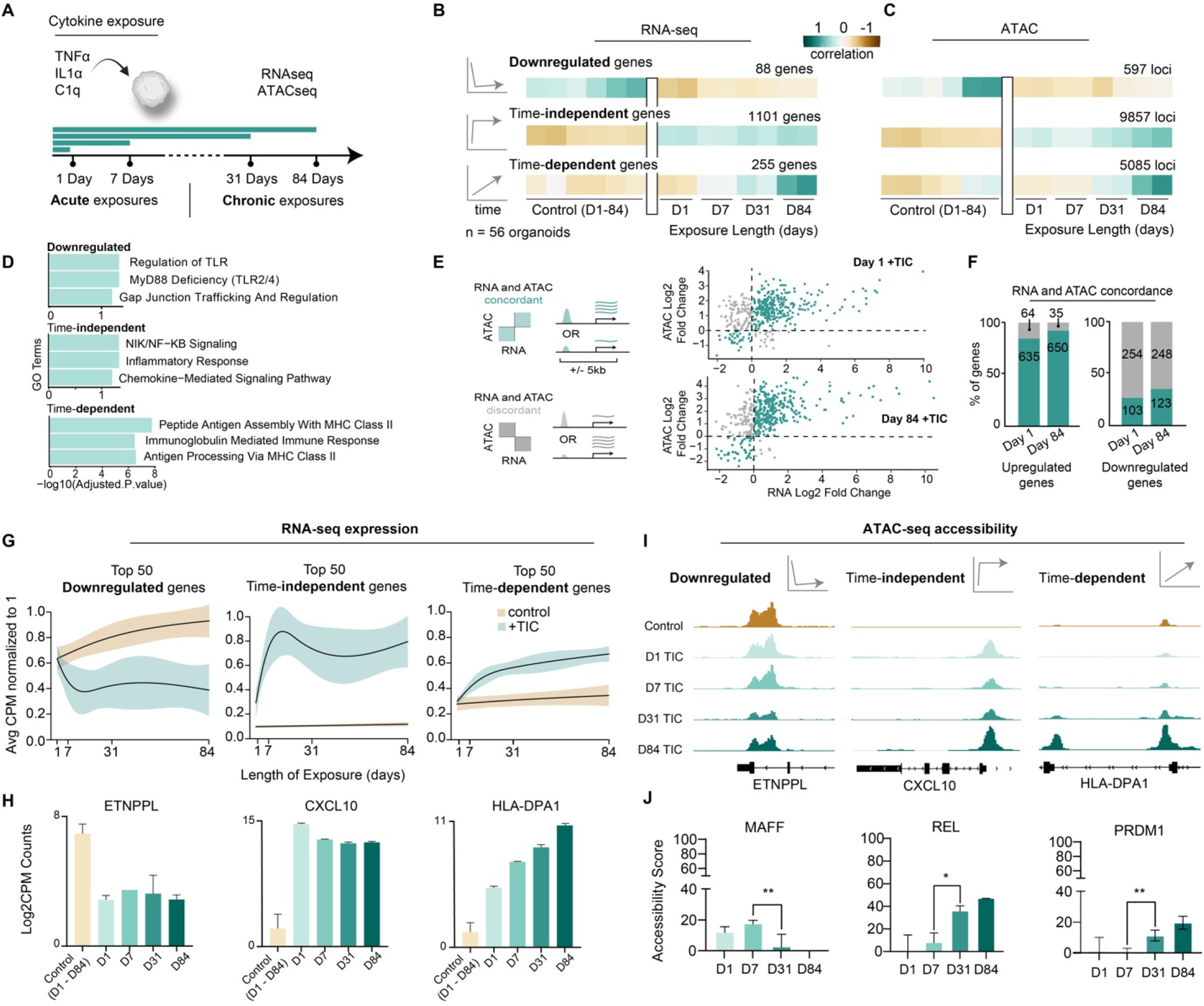
Astrocytes adopt distinct acute and chronic reactive states over time. **(A)** hCOs were exposed to TIC for 1, 7, 31, or 84 days. At each timepoint, astrocytes were purified and paired RNA- and ATAC-seq performed. **(B)** Heatmaps of selected WGCNA modules from both RNA-seq and ATAC-seq on purified control and reactive astrocytes at each timepoint. RNA parameters: minModuleSize=200, mergeCutHeight=0.15, ATAC parameters: minModuleSize=400, mergeCutHeight=0.15. **(C)** GO terms for genes in downregulated, time-independent, and time-dependent modules. **(D)** RNA and ATAC correlation scatter plots of DEGs and DARs plotted by log2FC. **(E)** RNA and ATAC-seq peaks were plotted together to assess correlation of genomic and transcriptomic responses over time after TIC exposure. **(F)** Quantification of RNA and ATAC correlation scatter plots. **(G)** Generalized additive model (GAM) plots of the top 50 variably expressed genes in each of the three selected WGCNA modules. Shaded regions correspond to the 95% CI. **(H)** Log2 CPM plotted for marker genes from each selected WGCNA module. **(I)** IGV plots of differentially accessible genomic loci found in the downregulated, time-independent, or time-dependent WGCNA modules. **(J)** TF bar charts: Accessibility scores were calculated using PECA2. The transcription factors shown have regulatory potentials that mirror the downregulated, time-independent, and time-dependent gene modules. MAFF: *p=0.0436, unpaired t-test, REL: **p=0.0091, unpaired t-test PRDM1: *p=0.013, unpaired t-test.

Gene ontology (GO) analysis identified terms associated with immune responses across all 3 modules, with Toll-like receptor signaling in the downregulated group, inflammatory response in the immediate-responding genes (time-independent), and surprisingly, antigen processing and presentation / major histocompatibility complex class II (MHCII) in the time-dependent module (**Fig. 2D**).

To assess whether inflammatory gene activation required prior chromatin priming, we examined concordance between differentially expressed genes and nearby differentially accessible regions (1,056 DARs, located within +/-5kb of each gene). For genes induced by TIC exposure (both acutely and chronically), we observed strong concordance between gene expression and chromatin accessibility (**Fig. 2E-F**). In contrast, we observed the opposite trend with genes downregulated upon TIC exposure, which exhibited largely discordant patterns between gene expression and chromatin accessibility (**Fig. 2E-F**), suggesting that these loci may remain poised for rapid reactivation once the inflammatory stimulus is removed. To visualize these distinct regulatory programs, we applied generalized additive models (GAMs) to the RNA and ATAC data (**Fig. 2G**) and highlighted representative gene/peak trajectories within each module (**Fig. 2H–I**).

We found that transcription factor regulatory activity varied substantially with the duration of inflammation. To identify candidate regulators driving these responses, we used PECA2, a computational tool that integrates ATAC- and RNA-seq data to infer transcription factors associated with each inflammatory gene module (**Fig. 2J, Supp. Data 4-7**). These identified transcription factors represent putative regulators of these gene modules. Together, these findings define a distinct chronic inflammatory astrocyte state, characterized by transcriptional and epigenetic features that diverge from the acutely reactive state.

### Reactive astrocytes can return to quiescence after both acute and chronic inflammatory conditions

Astrocytes respond uniquely to the duration of inflammatory exposure, but are these changes permanent? To assess the plasticity of the reactive state, we next tested the ability of inflammatory astrocytes to return to quiescence following withdrawal of the inflammatory stimulus. We exposed hCOs to TIC for either 7 (acute exposure) or 31 (chronic exposure) days, followed by removal of TIC and a subsequent withdrawal period (**Fig. 3A**). We collected hCOs at regular 1-week or ∼2-week intervals following the acute or chronic exposure periods to monitor the plasticity and/or permanence of the reactive signatures. At each timepoint, we dissociated hCOs, purified astrocytes, and performed paired ATAC- and RNA-seq. As expected, 7 days of acute TIC exposure induced a robust transcriptomic and chromatin accessibility signature of the time-independent inflammatory astrocyte module, but these changes returned almost entirely to baseline after 7 days of withdrawal (**Fig. 3B-C, F**). We observed a similar trend in the time-dependent module, although the initial induction of these genes and chromatin loci was less pronounced, consistent with their sensitivity to exposure duration (**Fig. 3D–F**). Remarkably, this plasticity persisted even following chronic exposure: both time-independent and time-dependent DEGs and DARs returned to baseline levels after 31 days of TIC treatment and subsequent withdrawal (**Fig. 3F–L**). In contrast, the downregulated module exhibited a slower return to quiescent levels following both acute and chronic exposure (**Supp. Fig. 2A, C**). Interestingly, while expression of these genes was suppressed, chromatin accessibility at the corresponding loci remained largely unchanged—highlighting a discordance between transcriptional downregulation and epigenomic remodeling (**Supp. Fig. 3B, D– E**)

**Fig. 3.**
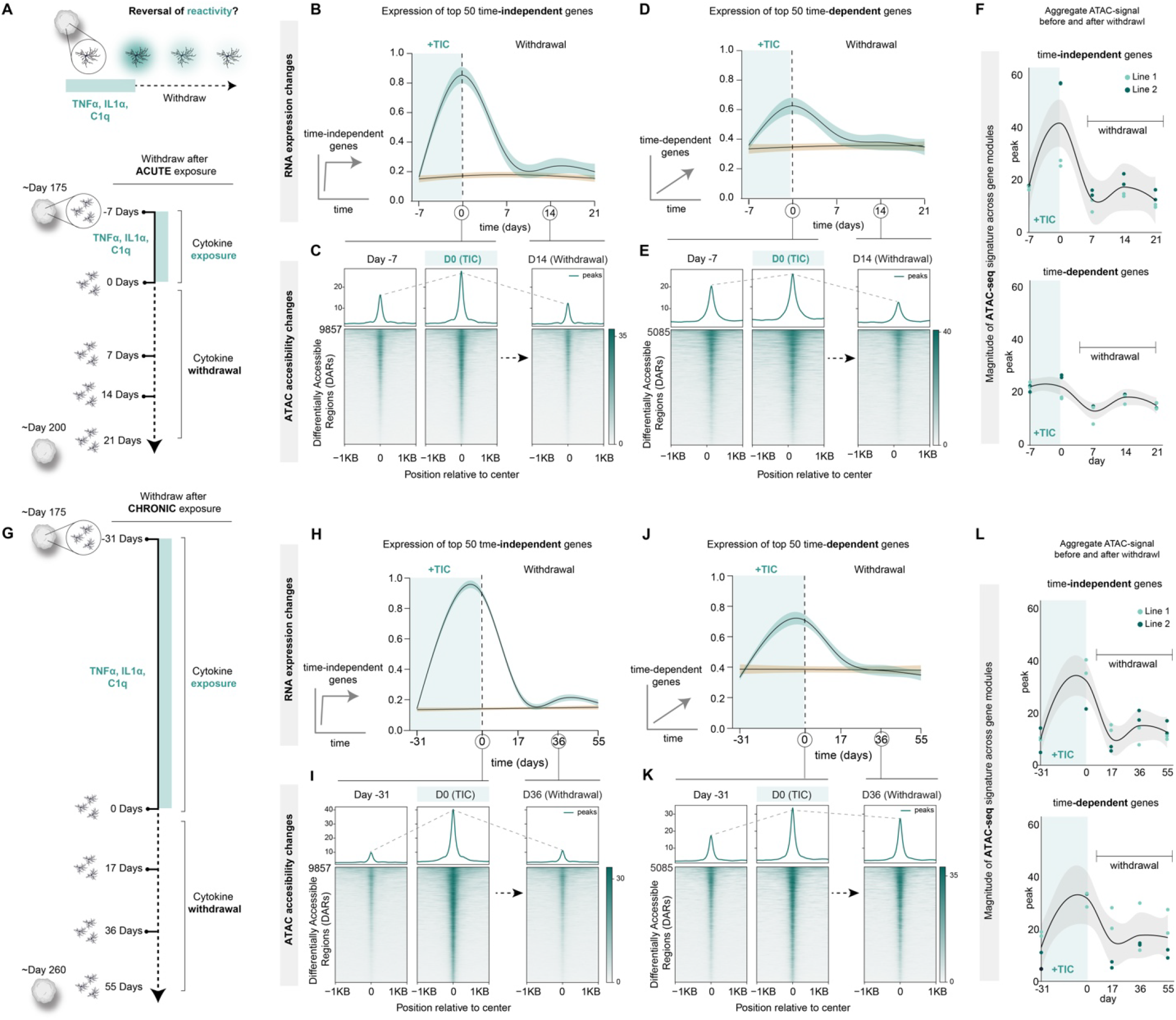
Resolution of astrocyte reactivity depends on duration of inflammatory exposure. **(A)** hCOs were exposed to TIC for 7 days, then withdrawn from TIC and cultured in normal media. At regular intervals, astrocytes were purified and paired RNA- and ATAC-seq performed. (n= 2 iPSC lines, ∼64 hCOs/line). **(B)** GAM plot depicting normalized Log2 CPMs of the top 50 most variably-expressed time-independent genes after acute TIC exposure and withdrawal. **(C)** Importance plots for control, 7 day +TIC exposure, and 21 days of withdrawal from TIC exposure for time-independent gene associated DARs after acute TIC exposure and withdrawal. DARs are aligned in the middle of the plot, with +/-2kb from the DAR. The most variably-accessible DARs are plotted toward the top. **(D)** GAM plot depicting normalized Log2 CPMs of the top 50 most variably-expressed time-dependent genes after acute TIC exposure and withdrawal. **(E)** Importance plots for control, 7 day +TIC exposure, and 21 days of withdrawal from TIC exposure for time-dependent gene-associated DARs after acute TIC exposure and withdrawal. **(F)** Magnitude of ATAC peaks within either the time-independent or time dependent gene modules across the acute withdrawal timeline. These points correspond to the importance plots in C and E. **(G)** hCOs were exposed to TIC for 31 days, then withdrawn from TIC and cultured in normal media. At regular intervals, astrocytes were purified and paired RNA- and ATAC-seq performed. (n= 2 iPSC lines, ∼64 hCOs/line). **(H)** GAM plot depicting normalized Log2 CPMs of the top 50 most variably-expressed time-independent genes after chronic TIC exposure and withdrawal. **(I)** Importance plots for control, 31 day +TIC exposure, and 67 days of withdrawal from TIC exposure for time-independent gene associated DARs after chronic TIC exposure and withdrawal. **(J)** GAM plot depicting normalized Log2 CPMs of the top 50 most variably-expressed time-dependent genes after acute TIC exposure and withdrawal. **(K)** Importance plots for control, 31 day +TIC exposure, and 67 days of withdrawal from TIC exposure for time-dependent gene associated DARs after chronic TIC exposure and withdrawal. **(L)** Magnitude of ATAC peaks within either the time-independent or time dependent gene modules across the chronic withdrawal timeline. These points correspond to the importance plots in I and K.

### Astrocytes have detectable MHCII protein after chronic inflammation

While the role of reactive astrocytes in inflammation is a growing focus in the field of glial biology, the involvement of MHCII in the astrocyte-mediated immune responses remains poorly understood. Prior studies have consistently reported upregulation of MHC class I molecules (primarily HLA-A – HLA-E alleles) in astrocytes following acute inflammatory stimuli, such as LPS injection in mice (7, 8) or TIC exposure *in vitro* (17–19). As expected, we observed robust induction of MHC class I genes in our dataset. Unexpectedly, we also found that chronic TIC exposure induced astrocytic expression of MHCII genes—components of professional antigen presentation machinery—despite astrocytes not being professional antigen-presenting cells (35, 36). In rodents, astrocytic MHCII expression has only been reported in rare contexts, typically in response to IFN-γ, which directly activates the MHCII transcriptional program (23, 37–39). Therefore, we sought to validate our preliminary findings that chronic inflammation (in the absence of IFN-γ) could also induce MHCII machinery in human astrocytes. First, to determine whether transcriptional upregulation of MHCII was accompanied by protein translation, we performed immunostaining on hCOs exposed to TIC for varying durations, ranging from 1 day to 3 months (**Fig. 4A-B**). At baseline, MHCII protein was virtually undetectable. However, by day 7, we observed localized immunofluorescence at the organoid periphery, likely where cytokine concentrations are highest. MHCII signal continued to intensify with prolonged TIC exposure. To confirm the cellular identity of MHCII-expressing cells, we next performed co-immunostaining for astrocyte markers, verifying that MHCII protein was indeed expressed within astrocytes (**Fig. 4C–D**), and not within a contaminating non-glial or non-canonical cell population.

**Fig. 4.**
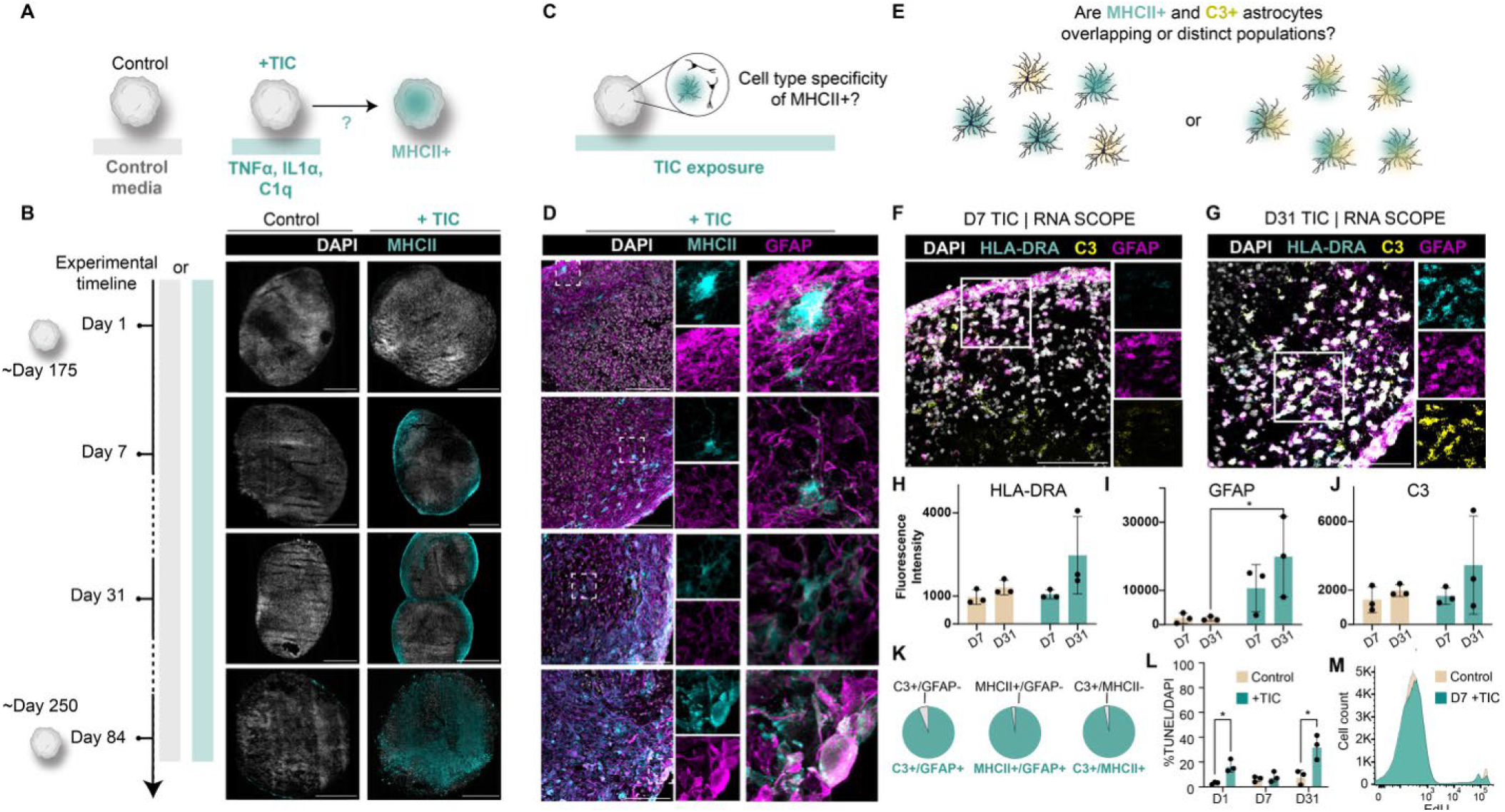
MHCII expression emerges in astrocytes with prolonged TIC exposure. **(A)** hCOs were cultured with either control media or +TIC media that was refreshed every 3 days for 1, 7,31, or 84 days. **(B)** hCOs sections immunostained for DAPI (grey) and MHCII (cyan). hCOs were either cultured in normal media or media +TIC for 1-84 days. Tiled images were taken at 40x magnification and stitched. Scale= 500µm, n= 2 iPSC lines. **(C)** hCOs were cultured +TIC to understand whether MHCII expression was specific to astrocytes **(D)** hCOs sections immunostained for DAPI (grey), MHCII (cyan), and GFAP (magenta). hCOs were cultured in media +TIC for 1-84 days. Single-channel (middle) and overlay (right) insets are demarked by white boxes (left panel). scale= 100µm, n=2 iPSC lines. (**E-G)** RNAscope image of hCOs exposed to either 7 or 31 days +TIC. Tissue sections were probed for HLA-DRA (MHCII allele, cyan), C3 (yellow), and GFAP (astrocytes, magenta). Single channel insets demarked by white boxes. Scale= 100µm, n = 1 iPSC line, 3 sections/condition. **(H-J)** Quantification of mean fluorescence intensity for HLA-DRA, C3, and GFAP in panel (C). Statistical test: 2-way ANOVA, p-value=0.012. n = 1 iPSC line, 2 hCS/condition. **(K)** Quantification of D31 RNAscope probe colocalization. Left: C3 and GFAP (94%), Middle: HLA-DRA and GFAP (100%), Right: C3 and HLA-DRA (97%). **(L)** Quantification of TUNEL stain on hCOs sections either cultured in normal media (control) or media +TIC for 1, 7, or 31 days. Statistical test: 2-way ANOVA, p-values: D1 TIC=0.0369, D31 TIC=0.0306. **(M)** Flow analysis of hCOs cultured in normal media + EdU or media +TIC +EdU for 7 days. n=3 hCOs/condition, 12 hCOs total.

### Astrocytes can express both early inflammatory genes and MHCII

To determine whether MHCII+ astrocytes represent a distinct population from those acutely-activated by inflammation (which express time-independent markers), we performed RNAscope using probes for HLA-DRA (an MHCII allele), C3 (a time-independent reactive marker), and GFAP at both 7 and 31 days of TIC exposure (**Fig. 4E-G**). As expected, we observed high levels of MHCII transcripts following 31 days of TIC exposure (**Fig. 4G, H**), consistent with MHCII’s classification as a time-dependent response. Additionally, transcript levels for both C3 and GFAP also increased in abundance from day 7 to day 31 of TIC exposure (**Fig. 4I-J**), indicating progressive astrocyte reactivity over time. Notably, by day 31, 100% of MHCII^+^cells colocalized with GFAP, and 97% colocalized with C3, suggesting that MHCII expression arises exclusively within astrocytes—and more specifically, within cells that likely had already adopted a reactive state earlier in the inflammatory timeline (**Fig. 4K**). These data support a model in which chronically stimulated astrocytes initially acquire a classical inflammatory phenotype and subsequently transition into a distinct MHCII-expressing state.

### Inflammatory conditions lead to an increase in cell death, but not a rise in proliferation

Previous studies suggest that inflammatory reactive astrocytes may lead to increased cell death. To test this in our system, we performed a TUNEL stain on hCOs exposed to TIC for either 1 or 31 days. TIC treatment led to a marked increase in cell death at both timepoints— 16.8% and 32.4% of DAPI^+^nuclei, respectively—compared to age-matched controls (3.9–8%) (**Fig. 4L**). The identity of dying cells, however, is difficult to define due to the nature of this assay. We next asked whether the emergence of an MHCII^+^astrocyte population could be explained by selective expansion of a rare, proliferative reactive astrocyte subtype. To test this, we co-exposed hCOs to TIC and EdU for either 1 or 7 days and assessed proliferation by flow cytometry. Surprisingly, we observed a slight reduction in EdU incorporation in TIC-treated hCOs compared to controls, suggesting there is not inflammation-induced proliferation of astrocytes or a specific reactive subpopulation in this model of reactivity (**Fig. 4M, Supp. Fig. 4A-B**).

### Reactive astrocytes present MHCII on the cell surface

Our MHCII immunostaining demonstrated progressive expression in the setting of TIC but lacks the ability to distinguish between surface-exposed, a hallmark of antigen presenting-competent MHCII complexes, and MHCII pools present in intracellular organelles. Therefore, to determine whether MHCII was localized to the astrocyte surface, an essential step for functional antigen presentation, we turned to an alternative model: organotypic slice cultures of fetal human cortical tissue. We selected this approach for two reasons: (1) fetal brain slices yield significantly more cells than hCOs, and (2) they contain microglia, which are absent in hCOs. To test whether astrocyte MHCII expression could be an artifact of microglia-deficient in vitro conditions, we vibratome sectioned GW18-21 human cortical tissue and cultured the slices on transwell inserts. We then exposed these sections to either IFN-γ (positive control) or TIC to induce MHCII expression (**Fig. 5A-B**). After dissociating the tissue sections into a single-cell suspension, we performed FACS to separate myeloid cells from astrocytes and then measured MHCII levels in each population (**Fig. 5C**). We noted similar levels of MHCII expression in both myeloid and astrocytic populations (**Supp. Fig. 5C**). In the absence of membrane permeabilization, FACS detection was limited to MHCII expressed on the cell surface. Strikingly, we were able to sort MHCII^+^astrocytes from both IFN-γ- and TIC-treated slices, confirming that MHCII reaches the astrocyte surface even in the presence of microglia (**Fig. 5D– F**). We then performed RNA-seq on the sorted MHCII^+^and MHCII^−^astrocyte populations. MHCII^+^astrocytes showed a marked upregulation of MHCII genes as well as the key MHCII regulator CIITA (**Fig. 5G–H**).

**Fig. 5.**
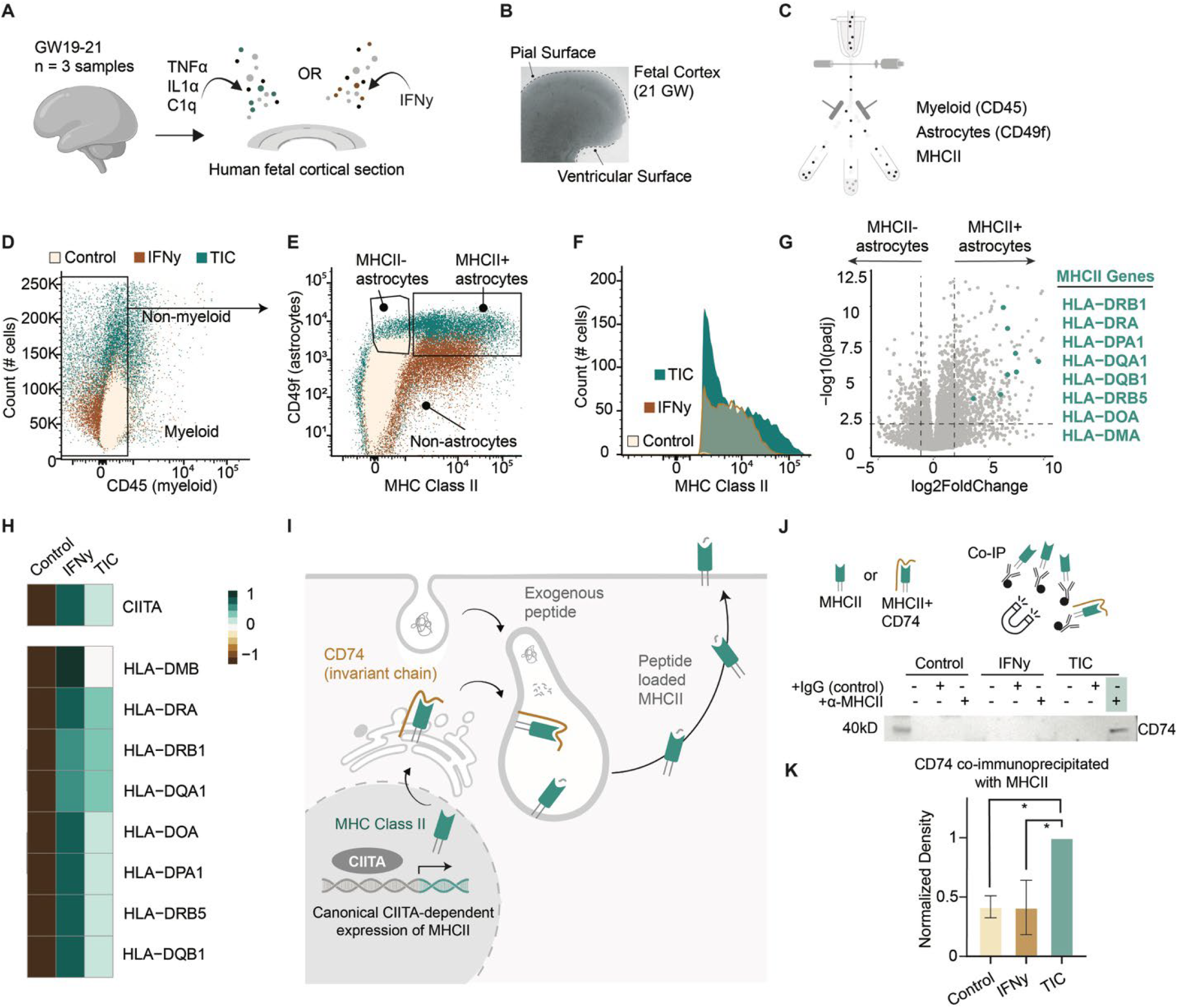
TIC induces surface MHCII expression in fetal human astrocytes. **(A)** 300µm GW18-21 fetal cortical sections were cultured on transwell inserts for 7 days either with no additives, +IFN-γ (2 days), or +TIC (7 days). n = 3 samples. **(B)** Phase image of 21GW fetal human cortical section, with pial and ventricular surfaces denoted. **(C)** FACS sorting scheme. **(D)** Dot plot showing negative selection against myeloid cells during FACS. **(E)** Dot plot showing positive selection of MHCII+ CD49f+ cells. **(F)** Histogram of MHCII+ CD49f+ astrocytes across all 3 conditions. **(G)** Volcano plot of RNA-seq showing MHCII genes are differentially expressed in fetal astrocytes after TIC exposure. **(H)** Heatmap of control, +IFN-γ, and +TIC MHCII mRNA expression **(I)** Schematic of antigen processing and presentation via MHCII and CD74 **(J)** Western blot of CD74 protein co-immunoprecipitated with MHCII from fetal human astrocytes. **(K)** Quantification of Western blot in (J). 1-way ANOVA: TIC vs control p-value=0.018, TIC vs IFNγ p-value=0.011

### MHCII complexes with functional molecules in reactive astrocytes, such as CD74, the invariant chain

To further explore MHCII functionality in reactive astrocytes, we immunopanned astrocytes from GW18-21 human cortex and cultured in either standard neurobasal media or neurobasal media supplemented with IFN-y (for 2 days) or TIC (for 7 days) to induce putative MHCII expression. We next lysed the cells and performed immunoprecipitation (IP) for MHCII and probed for MHCII-associated proteins. We found the invariant chain, CD74, immunoprecipitated with the MHCII complex in astrocytes exposed to TIC compared to unexposed astrocytes (p-value=0.018) and IFN-y exposed astrocytes (p-value= 0.011) (**Fig. 5I-K, Supp. Fig. 6**).

### MHCII-bound peptides can be identified via LC-MS/MS

To identify proteins that associate with MHCII chains and distinguish them from putative peptides presented by MHCII as antigens, we performed liquid chromatography-tandem mass spectrometry (LC-MS/MS) on proteins and peptides coimmunoprecipitated with MHCII (**Fig. 6A**). We confirmed the precipitation of MHCII complexes by the presence of MHCII and CD74 in the IP samples (**Fig. 6B, Supp. Data 8**). We next sought to distinguish between proteins that associate with MHCII and those that are presented as antigens. We classified proteins consistently enriched across all IP samples as likely MHCII-associated or complexed proteins (e.g., HLA-DPB1, HLA-DRB1, CD74) (**Fig. 6C**). In contrast, we considered peptides enriched only in a subset of IP samples, but highly abundant within them, as likely MHCII-presented peptides (**Fig. 6D**). We further categorized these putative peptides by analyzing the peptide sequences for trypsin cleavage sites. Because peptides were digested with trypsin prior to LC-MS/MS, we reasoned that trypsin cleavage patterns could differentiate between associated full-length proteins and shorter presented peptides. We hypothesized that peptides containing trypsin cleavage sites at both ends are more likely derived from intact proteins associated with MHCII, whereas putative peptides would likely be shorter (up to ∼30 amino acids), and may only possess a single trypsin cleavage site, having been initially processed by endolysosomal cathepsins with a different preferred cut consensus site (37, 38). Supporting this distinction, we found that 84% of putative MHCII-associated peptides possessed bilateral trypsin cleavage sites, compared to only 44% of putative MHCII-presented peptides (**Fig. 6E**).

**Fig. 6.**
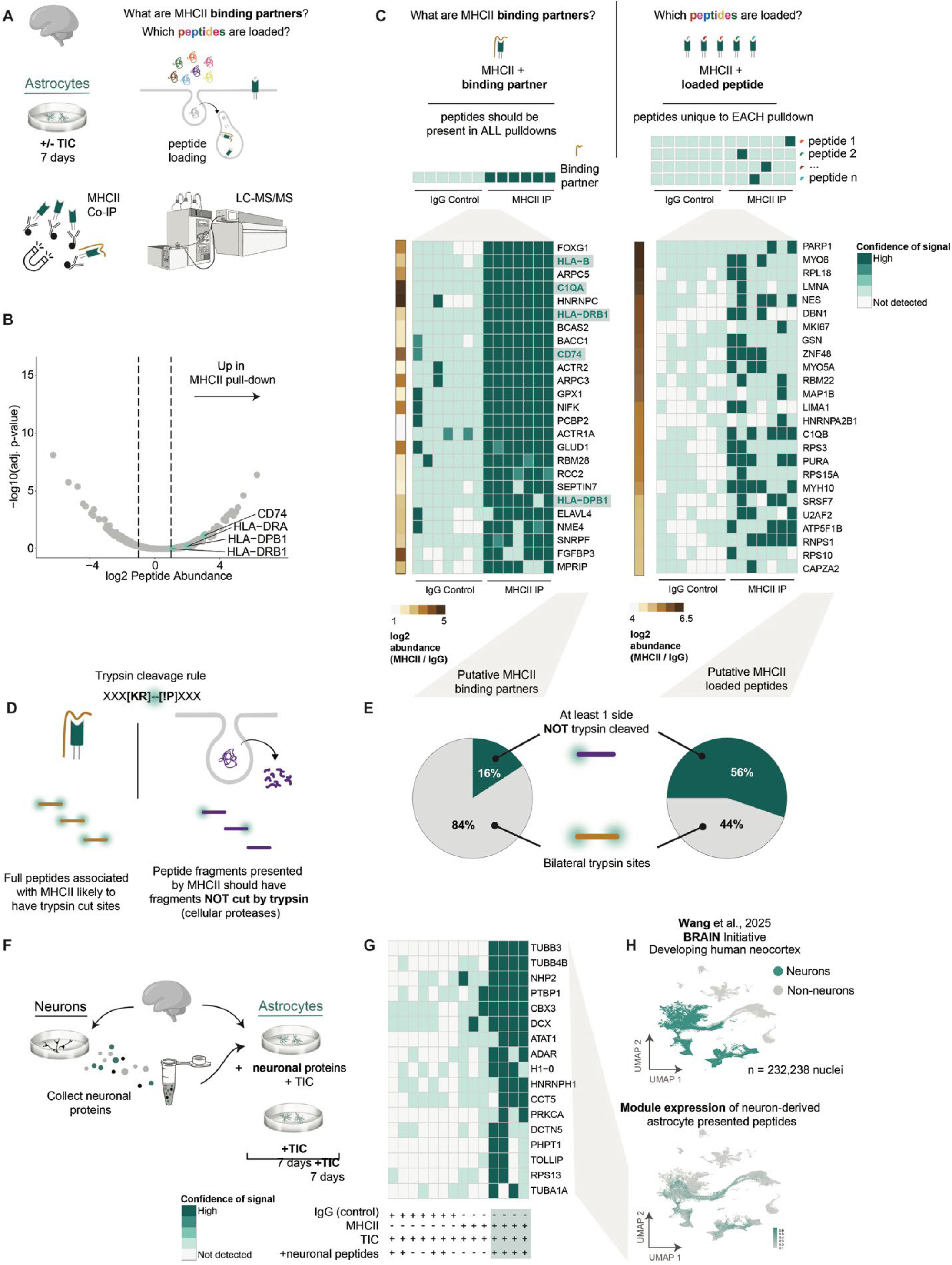
Fetal astrocytes acquire functional antigen-presenting features via MHCII. **(A)** GW18-21 astrocytes isolated from fetal human cortices were cultured -/+TIC for 7 days and peptides bound to or presented by MHCII were immunoprecipitated and identified via LC-MS/MS. **(B)** Peptides found in all MHCII IP samples but not the IgG controls were considered to be proteins that associate with MHCII throughout processing and presentation. Highly abundant peptides that were found in only a subset of MHCII IP samples but not IgG controls were considered putative presented peptides. **(C)** Heatmap depicting confidence of mass spectrometry signal and log2 peptide abundance for the top 25 peptides enriched across all MHCII IP samples. **(D)** Heatmap depicting confidence of mass spectrometry signal and log2 peptide abundance for the top 25 most abundant peptides found enriched in at least one MHCII IP sample. **(E)** Neurons were isolated from GW18-21 human fetal cortices and total cell protein fractions were collected. Astrocytes were isolated from GW18-21 human fetal cortices and cultured either unstimulated, +TIC, or +TIC and neuronal proteins for 7 days. Proteins immunoprecipitated with MHCII or IgG control were identified via LC-MS/MS. **(F)** Prospective antigens presented by MHCII should have one or fewer ends of the peptide sequence featuring a trypsin cleavage site. **(G)** Heatmap depicting confidence of mass spectrometry signal for the top 25 most abundant peptides with only one end cleaved by trypsin. **(H)** Left: UMAP clustering of snRNA-seq of the developing human cortex (Wang et al., 2025). Right: gene expression overlay of the proteins found in **(G)**.

To explore what peptides astrocytes might present under physiological conditions, we considered that during human brain development, neuronal turnover is high and apoptotic debris is abundant. We hypothesized that reactive astrocytes could process and present peptides derived from neighboring dying neurons. To test this, we isolated neurons and astrocytes from human fetal cortical samples using immunopanning. We induced astrocyte MHCII expression by exposing them to TIC cytokines. Simultaneously, we lysed the isolated neurons into a protein slurry and added this lysate to the reactive astrocyte cultures for one week (**Fig. 6F**). After exposure, we immunoprecipitated MHCII-bound peptides (along with IgG controls from matched cultures) and performed LC-MS/MS. We specifically searched for peptides that were enriched in the MHCII IP samples but absent from IgG controls, including IgG pulldowns from cultures exposed to neuronal peptides. We further filtered this list for peptides that lacked bilateral trypsin cleavage sites, indicative of endolysosomal processed antigens rather than full-length co-precipitating proteins. Strikingly, we identified many abundant MHCII-associated peptides corresponding to canonical neuronal proteins, including TUBB3, TUBB4B, and DCX (**Fig. 6G**). To further verify their neuronal origin, we mapped the expression of our top 25 proteins onto a single-nucleus RNA-seq dataset from the developing human neocortex (Wang et al.), with minimal or absent expression in non-neuronal clusters (**Fig. 6H**). This pattern underscores that these peptides are not likely endogenously expressed by astrocytes but instead derive from exogenous neuronal sources. Together, these results strongly suggest that reactive astrocytes have the capacity to process and present neuronal peptides on MHCII.

## Discussion

### Immature astrocytes can become reactive

A central question we sought to address was whether prenatal, immature astrocytes, like those in hCOs, can mount a reactive response. Much of our current understanding of astrocyte reactivity is based on studies conducted in murine models, particularly in mature or aged animals. However, recent work using 2D hiPSC-derived cultures supports the idea that immature astrocytes can exhibit inflammatory phenotypes in response to cytokine exposure (Barbar et al. 2020; M. D. Smith et al. 2022). Our observation of a robust inflammatory response in 3D hCOs extends these findings and highlights the utility of organoids as a reproducible, tractable system for modeling human astrocyte reactivity. Notably, we directly compared inflammatory responses in hCO-derived astrocytes to primary human fetal astrocytes and found strong concordance in transcriptional profiles. Interestingly, fetal astrocytes displayed a less extensive transcriptional response, with fewer genes upregulated following TIC exposure than their hCO-derived counterparts.

An important question that remains is what functional consequences arise from the reactivity of fetal astrocytes. Numerous prenatal and perinatal conditions (hypoxic ischemic encephalopathy, perinatal infection, placental abruption) are associated with neurological injury, yet the extent to which astrocyte reactivity contributes to their pathogenesis remains unclear (39–41). Understanding whether fetal astrocyte responses exacerbate or mitigate damage in these contexts will be an important direction for future research. Furthermore, comparing inflammatory signatures in immature astrocytes to those observed in neurodegenerative diseases could offer insight into how astrocyte function evolves across the lifespan and disease states.

Our findings suggest that the hCO model, despite being a reductionist system lacking pathogenic mutations, recapitulates key reactive astrocyte signatures observed across a range of neurological diseases and injury states, both in murine models and in postmortem sn-RNAseq of human tissue (10–12,20). For the field, defining the extent of astrocyte plasticity is a critical first step toward understanding how these cells might be targeted to improve outcomes in disease or injury. Notably, our demonstration that even chronically activated reactive astrocytes can return to a quiescent state following withdrawal of inflammatory stimuli supports the idea that, in vivo, modulating inflammation may be a viable therapeutic strategy for mitigating long-term neurological dysfunction.

### Reactive astrocytes possess an extremely plastic genomic and transcriptomic landscape

Previous studies have shown that inflammatory astrocytes can regain pre-injury transcriptional signatures following acute stimuli (14, 15). Here, we extended this question to chronic inflammation: do astrocytes retain the capacity to return to a quiescent state after prolonged activation, or do they remain transcriptionally “scarred”? This has important translational implications. First, most neurological injuries and diseases involve sustained inflammation rather than acute insults (22, 42). Second, our findings suggest that removing pro-inflammatory signals may be sufficient to reverse astrocyte reactivity, thereby limiting secondary damage. Using paired RNA-seq and ATAC-seq, we show that astrocytes exposed to chronic TIC return to baseline levels of chromatin accessibility and gene expression, including inflammatory mediators and MHCII pathway components, following cytokine withdrawal. In vivo, such a reversal likely depends on reductions in inflammatory cues from neighboring CNS cell types. Importantly, astrocyte plasticity is not uniform and may be influenced by intrinsic factors such as genotype and brain region, especially in the context of neurodevelopmental or neurodegenerative disorders.

### Why do chronic reactive astrocytes upregulate MHCII?

One of the most unexpected findings from this study is that astrocytes exposed to chronic inflammation acquire antigen presentation capacity via upregulation of MHC class II (MHCII) molecules. This observation aligns with previous studies showing that genetic attenuation of astrocyte reactivity impairs CNS immune responses and exacerbates infection or inflammation (43–45). Moreover, recent data from iAstrocytes derived from multiple sclerosis (MS) patients revealed upregulation of HLA-DRA, an MHCII allele, further supporting the idea that astrocytic MHCII may play a meaningful role in neurological disorders (46). While MHCII activation has been well-characterized in professional antigen presenting cells (APCs) like microglia, its regulation in astrocytes remains poorly understood. In APCs, MHCII expression is driven by the non–DNA-binding coactivator CIITA, which must be recruited to the MHCII locus by additional transcription factors (47, 48). Whether astrocytes use the same regulatory machinery is unknown, and differences in these upstream pathways could explain the delayed MHCII induction observed in our model (49). Notably, while TIC-treated astrocytes do upregulate CIITA, the magnitude of induction is significantly lower than in IFN-γ–stimulated astrocytes.

What functional role MHCII plays in astrocytes, and whether it contributes to initiating, sustaining, or resolving neuroinflammation, remains an open and important question. Initially, we considered whether MHCII upregulation in hCOs might represent a compensatory artifact caused by the absence of a myeloid compartment. However, our experiments using human fetal cortical slices, which do contain microglia, argue against this interpretation. Even in the presence of professional antigen-presenting cells, astrocytes acquired MHCII expression and surface presentation, supporting the idea that this is a bona-fide astrocytic response to chronic inflammation. Other work has shown that OPCs are also capable of expressing MHCII in disease settings (11, 50), suggesting that non-microglial antigen presentation may be more widespread than previously appreciated.

FACS and co-immunoprecipitation confirmed that MHCII is surface-expressed and associates with CD74 in TIC-treated astrocytes, including those derived from fetal tissue, indicating functional trafficking and ruling out organoid-specific effects. Importantly, mass spectrometry also revealed a broad array of peptides potentially presented by astrocytic MHCII. The origin of these peptides likely includes some amount of exogenous material that astrocytes have endocytosed and processed. Supporting this, when we added neuronal proteins to our reactive astrocyte cultures, neuronal tubulins—abundant neuronal structural proteins—were among the most commonly identified peptides. These findings raise the possibility that astrocytes may engage in antigen uptake (perhaps via their phagocytic capacity) under chronic inflammatory conditions.

### What is the role of antigen presentation in astrocytes?

Together, our findings suggest that astrocytes are not merely passive expressers of MHCII but may actively participate in antigen processing and presentation. However, the biological consequences of this remain unclear. Do MHCII^+^astrocytes interact with T cells in vivo? Do they shape local immune tone, promote tolerance, or contribute to pathology? Determining what astrocytes are presenting, to whom, and to what effect remains a key question for the field.

One yet-to-be-tested hypothesis is that astrocytic antigen presentation may serve an anti-inflammatory role (particularly in chronic settings) by promoting immune dampening rather than activation. Supporting this idea, we did not detect robust expression of CD80, a canonical co-stimulatory molecule required for full T cell activation, in MHCII^+^astrocytes. Since MHCII can present intracellular (self-derived) peptides (38), it is possible that, after prolonged inflammation, astrocytes may promote immune tolerance by presenting self-antigen in the absence of co-stimulatory signals (51).

This concept aligns with prior studies showing that reactive astrocytes can restrict inflammation through physical and molecular barriers—for example, by forming glial scars to contain immune cell infiltration following stroke or spinal cord injury (52–55). In this context, the delayed kinetics of MHCII induction in response to TIC may reflect a threshold-based mechanism, where astrocytes require sustained inflammatory signaling, or activation of multiple parallel pathways, before adopting an antigen-presenting phenotype that may ultimately help resolve inflammation.

## Methods

### Human cortical organoid cultures

Human cortical organoids were formed from three human induced pluripotent stem cell (hiPSC) lines (8858.3, 1363.1, and C4.1) (Paşca et al. 2015) following a previously published protocol. All lines were genotyped by SNP-array to confirm genomic integrity and regularly screened for mycoplasma. iPSC colonies at 80–90% confluency were detached from culture plates using Accutase and were formed into 3D spheroids using AggreWell^™^ plates. Following 3D formation, spheroids were treated daily in neural induction media (DMEM/F12 (Thermo Fisher Scientific, Cat. 11330057), Knockout Serum Replacement (Thermo Fisher Scientific, Cat. 10828028), MEM Non-Essential Amino Acids Solution (Thermo Fisher Scientific, Cat. 11140050), Glutamax (Thermo Fisher Scientific, Cat. 35050061), Pen/Strep (VWR, Cat. 16777), Beta-mercaptoethanol) supplemented with Dorsomorphin (Sigma-Aldrich, Cat. P5499–25MG, 5 μM) and SB-431542 (Selleck Chemicals, Cat. S1067, 10 μM) for 6 days. Following this treatment, organoids were treated daily with neural media (Neurobasal-A (Thermo Fisher Scientific, Cat. 10888022), B-27 Supplement minus vitamin A (Thermo Fisher Scientific, Cat. 12587010), Glutamax (Thermo Fisher Scientific, Cat. 35050061), Pen/Strep (VWR, Cat. 16777)) supplemented with EGF (R&D Systems, Cat. 236-EG-01M, 20 ng/mL) and FGF2 (R&D Systems, Cat. 233-FB-01M, 20 ng/mL) for 10 days, and every other day for days 16–24. At day 25, organoids were treated every other day with neural media supplemented with BDNF (PeproTech, Cat. 450-02-1mg, 20 ng/mL) and NT-3 (R&D Systems, Cat. 267-N3–005/CF, 20 ng/mL) to promote differentiation of progenitors. From day 43 onwards, organoids were fed every 3 days with neural media only.

To induce astrocyte reactivity in hCOs, TNF-α (30ng/mL, Cell Signaling Technology, 16769S), IL-1α (3ng/mL,Sigma-Aldrich, I2778), and C1q (400ng/mL, EMD Millipore, 204876) were added to neural media every 3 days.

### Tissue Dissociation

We used a previously established protocol (Zhang et al., 2016) to dissociate human cortical tissue, both from human cortical organoids and primary human fetal cortical tissue (GW17-21). All human tissue samples were obtained in compliance with policies outlined by the Emory School of Medicine IRB office. In short, we mechanically and enzymatically dissociated tissue using Papain (Worthington Biochemical, LS003126) at 20U/mL at 34^°^C for 1 hour, quenched with ovomucoid solution (Worthington Biochemical, LS003086), and triturated to obtain single-cell suspension. Once we had a single-cell suspension, we proceeded to purify individual cell types through immunopanning.

### Immunopanning

The single-cell suspension was added to a series of Petri dishes that were pre-coated with cell-type specific primary antibodies and incubated between 10-15 minutes. We transferred unbound cells to the next plate and rinsed the plate with bound cells with PBS until only bound cells remain. The antibodies we used include: a-CD45 (microglia/macrophages, R&D Systems, MAB1430), a-CD24 (neurons, Miltenyi Biotec, 130-108-037),and a-HepaCAM (astrocytes, R&D Systems, MAB4108). We then incubated cells in a trypsin solution (Sigma-Aldrich, T9935-100MG) at 37^°^C for 10 minutes, quenched with 30% FBS (MedSupply Partners, 62-1300-1HI), and gently dislodged cells from the plates for culturing.

For cell culture, we pelleted cells, then plated them on poly-D-lysine-coated plastic coverslips in a Neurobasal-based serum-free medium supplemented with Antibiotic-Antimycotic (Gibco, 15240062) (composition in Supplemental Information). We performed complete media changes after 24 hours and then every 3 days after for up to approximately 2 weeks.

### RNA extraction

Total RNA was extracted from cells using the RNeasy Micro kit (QIAGEN, 74004) according to the manufacturer’s protocol. We assessed the quality of the RNA using the Bioanalyzer (Agilent, Eukaryotic Total RNA Nano or Pico kits). RNA samples with an RNA integrity number (RIN) of less than 9 were discarded.

### cDNA Library preparation

We created bulk RNA-seq libraries using the SMART-Seq^®^ v4 Ultra^®^ Low Input RNA Kit for Sequencing (Takara Bio, 634891) according to the manufacturer’s protocol to perform end repair, adaptor ligation, and 11 cycles of PCR amplification. We assessed library quality via Bioanalyzer (Agilent, High Sensitivity DNA kit). We then sequenced libraries on Illumina HiSeq 2500 or NovaSeqX Plus instrument to a depth of at least 20 million paired-end reads per sample.

### RNA-seq Analysis

Fastq files were processed as follows: files were trimmed using Trimmomatic, aligned to hg19 with STAR, and reads summarized with featureCounts. Samples that had an alignment rate of higher than 80% were used in downstream processing in Rstudio. Reads were normalized and differential expression analysis was performed with DESeq2. Genes expressed in similar patterns were sorted into modules using the WGCNA package.

### ATAC-seq Library Preparation

Bulk ATAC-seq libraries were prepared following the previously established Omni-ATAC protocol (56) with minor modifications (57). Briefly, astrocytes that were trypsinized during immunopanning were counted and 10,000-50,000 cells were washed with cold ATAC-seq resuspension buffer (RSB; 10 mM Tris-HCl pH 7.4, 10 mM NaCl, and 3 mM MgCl2 in water), and permeabilized with ATAC-seq lysis buffer (RSB supplemented with 0.1% NP40, 0.1% Tween-20, and 0.01% digitonin). For the transposition step, nuclei were resuspended in 44 µl of transposition mix (25 µl 2× TD buffer, 2.5 µl transposase, 16.5 µl PBS, 0.5 µl 1% digitonin, and 0.5 µl 10% Tween-20) and incubated at 37^°^ C for 40 min in a thermomixer with shaking at 1,000 RPM. The DNA fragments were cleaned up with the DNA Clean & Concentrator kit (Zymo Research, D4014) and PCR amplified for 8-12 cycles with Illumina Nextera adaptors using Kapa SYBR^®^ FAST (Roche, 07959427001). While the PCR was running, we watched for a change in fluorescence of 100,000 or a ΔRn of 0.1, after which PCR was stopped and the samples removed. After amplification, a second cleanup with 1.8x Ampure XP beads (Aline biosciences, Cat. C-1003-50) was performed and library eluted in 10mM Tris-HCl. Fragment size distribution and concentration were evaluated via Bioanalyzer High Sensitivity DNA kit (Agilent, 5067-4626), and libraries were sequenced using 2×150-bp reads on an Illumina HiSeq 2500 or NovaSeqX Plus instrument at a targeted depth of at least 50 million paired-end reads per sample.

### ATAC-seq Analysis

Reads were checked for quality using FastQC and FastQ screen. Reads were then trimmed using Trim Galore!, aligned to hg19 with Bowtie2, and files that had an alignment rate of higher than 80% were used. Mitochondrial reads, duplicates, non-unique alignment reads, and black-listed reads were removed using Samtools, PicardTools, and Bedtools. The final .bam files were then indexed and imported into RStudio for peak calling using the ChrAccR package and MACS2.

### Single Cell Prep with CMOs

We dissociated organoids into a single cell suspension using previously published methods (Zhang et al., 2016). Cells were incubated with CMOs according to protocols for Chromium Next-GEM single cell 3’ reagent kit v3.1 (dual-index) with cell multiplexing (PN-1000261 and PN-1000262).

We stained cells with 50nM calcein-AM and 4µM ethidium homidimer-1 from the LIVE/DEAD^™^ Viability/Cytotoxicity Kit (Thermofisher Scientific, L3224) prior to cell sorting. Live (calcein^+^, ethidium-) cells were sorted on a BD FACS Aria II. Cells were sorted into 4% BSA, supplemented with Protector RNAse inhibitor (1:1000, Sigma 03335402001) and transcriptional inhibitor Actinomycin D (1:1000, Sigma-Aldrich, A1410).

Actinomycin D was reconstituted in DMSO at stock concentration of 5mg/ml and aliquoted and stored at -20^°^C protected from light for up to 1 month.

Sequencing was performed by Admera Health on a NovaSeq S4 2×150 to achieve 50-100k reads per cell.

### 10X Analysis

Cells were demultiplexed using Cell Ranger. Analysis was performed in RStudio using Seurat. For quality control, cells with <10% mitochondrial genes were excluded. Cells containing nFeature_RNA>500 features were included for analysis.

### Immunofluorescence

Whole hCOs were fixed in 4% PFA (in 0.2M phosphate buffer) for 1-24hrs overnight at 4C (or 3hrs room temp) depending on organoid age/size. hCOs were cryopreserved in 30% sucrose in 1xPBS for 24-48 hours until no longer floating, before embedding in OCT (VWR 25608930) and stored at -80C. Cryosections of 12-16 µM were mounted on Superfrost Plus glass slides (Thermofisher, 12-550-15), allowed to anneal at RT, then stored at -80C.

5% normal donkey serum (Jackson ImmunoResearch, 017-000-1219) with 0.15% Triton X-100 was used for blocking and staining. Primary antibodies were incubated overnight at 4C, and secondary antibodies were incubated 1hr at room temperature. Hoechst (1:2000) or DAPI (Thermo Fisher D3571, 300nM/1:1000) were used for nuclear staining. Slides were mounted with coverslips and Fluoromount-G (Thermo Fisher, 00-4958-02) and cured overnight at RT.

### Imaging

Slides were imaged on a Leica Stellaris 5 confocal microscope using standard procedures.

Antibodies List:

GFAP, Agilent Dako, Z0334, 1:1000 GFAP, BioLegend, PCK-591P, 1:1000 MAP2, Abcam, ab11267, 1:1000

MHCII, NeoBiotechnologies (Thermo), 2967R, 1:200

### EdU Staining for FACS

hCOs were fed with neural media containing 10 µM EdU and TIC or only 10 µM EdU for either 24 hours or 7 days (replenished every 3 days). hCOs were dissociated according to our tissue dissociation protocol (see above), then cells were fixed and processed in accordance with the Click-iT Plus EdU Alexa Fluor 647 Flow Cytometry Kit instructions.

### Fluorescence-activated cell sorting

hCOs or fetal tissue sections were dissociated into single-cell suspensions according to the tissue dissociation protocol above. Cells were blocked in 1% BSA for 20 minutes then cells were counted and set aside for single-stain controls. Conjugated antibodies and Cytopainter Live cell stain were added to cells in dark at RT for 30 minutes. Antibodies were diluted with sterile FACS buffer (1x PBS w/o Ca/Mg2^+^, 1mM EDTA, 25mM HEPES pH 7.0, 1% heat-inactivated FBS), centrifuged, and resuspended in fresh FACS buffer.

Cells were sorted and collected using a FACS AriaII cell sorter into FACS buffer, centrifuged, then lysed for RNA isolation and downstream library creation and RNA-seq.

### FACS antibodies

MHCII, NeoBiotechnologies (Thermo), 2967R, 1:200, conjugated to AF594

CD49f-FITC, BD Biosciences, 555735, 1:200

CD45-BV711, BD Biosciences, 564358, 1:200

Live cell labeling kit-blue fluorescence, Abcam, ab187963, 1:1000

### Co-immunoprecipitation and Immunoblot

Primary cultured cells were scraped in protein extraction buffer (20mM Tris-HCl pH 7.5, 150mM NaCl, 5mM MgCl2, 1% NP-40, MilliQ water), briefly centrifuged, pulse sonicated at amplitude 20 on ice, snap frozen, and stored at -80C. Protein lysate concentration was determined by BCA assay (Thermo Scientific, 23227). 2mg of protein was aliquoted and 3ug of either an IgG (R&D Systems, AB-105-C) negative control antibody or MHCII (NeoBiotechnologies, RBM1-2967-P1) antibody was added to the protein and incubated at 4C overnight, with spinning.

Protein-antibody mixture was then incubated with Dynabeads (Thermo, 10006D) for 4 hours at RT with agitation. The beads were washed on a magnetic tube rack and eluted according to manufacturer’s instruction.

After elution from beads, protein was loaded onto an 8% Bis-Tris gel (Thermo, NW00080BOX) and run at 100V until the dye front reached the end of the gel. The gel was then removed from the cassette and blotted (Thermo, IB23001×3) using the iBlot system. The blot was blocked for 1 hour in 5% milk + 0.1% Tween20, then incubated in primary antibody overnight at 4C. The next day, the blot was washed in 1x PBS, incubated in secondary antibody for 1 hour, and washed again in 1x PBS. The blot was then incubated in chemiluminescent substrate (Thermo, 34577) for 5min at RT and imaged using the iBright 1500 Imaging System.

### Human fetal astrocyte culture with TIC and neuronal lysate

Astrocytes were purified via immunopanning and cultured in 2D as described above. Neurons were simultaneously purified via immunopanning (anti-CD24) and 20 million cells were collected and lysed in 1mL 1x PBS following 7 days of culture. This lysate was pulse sonicated at amplitude 20 on ice, snap frozen, and stored at -80C. Lysate protein quantification was calculated using a BCA assay and equal levels of 70µg were added to astrocyte culture plates every 3 days in the presence of TIC cytokines. After one week of culture, the cultures were collected for co-immunoprecipitation of MHCII as described above.

### Mass Spectrometry

Proteomic analyses were performed at the Proteomics Core of the Roy J. Carver Biotechnology Center at the University of Illinois Urbana-Champaign. Proteins were reduced and alkylated before being digested off the beads with LysC and trypsin overnight. On the following day, the digested peptides were desalted using StageTips. The peptides were then analyzed using a Q Exactive HF-X mass spectrometer (ThermoFisher Scientific) coupled to an UltiMate 3000 UHPLC (ThermoFisher Scientific) over a 45-minute reversed-phase gradient. The raw LC-MS data was searched against the Uniprot Homo sapiens reference proteome using Byonic v5.6 (Protein Metrics) implemented in ProteomeDiscoverer v2.4 (Thermo). Precursor ion-based label-free quantitation was accomplished with the Minora feature finder and Precursor ion quantifier nodes in ProteomeDiscoverer.

## Data and code availability

All datasets will be made available through the Gene Expression Omnibus (GEO). All scripts generated will be available without restrictions upon request at the Sloan Lab Github.

## Acknowledgements

This research project was supported in part by the Emory University School of Medicine Flow Cytometry Core and the Carver Proteomics Core at the University of Illinois at Urbana-Champaign.

## Funding

NIMH R01 MH125956 (SAS)

NINDS R01 NS123562 (SAS)

Sontag Foundation Distinguished Scholars Award (SAS)

NIH F32 ES031827 (HLW)

R01 ES034796 (VF)

Cure Alzheimer’s (VF)

## Author contributions

Conceptualization: EJH, SAS

Methodology: EJH, CS, MMS, HLW, AK, AS, BOH

Investigation: EJH

Visualization: EJH, MMS

Funding acquisition: SAS

Project administration: EJH, SAS

Supervision: SAS, VF

Writing – original draft: EJH, SAS

Writing – review & editing: EJH, SAS, MMS, VF

