## Supplemental Figures 1-6 for "Temporal regulation of human reactive astrocytes reveals their capacity for antigen presentation"

**Fig. S1.**

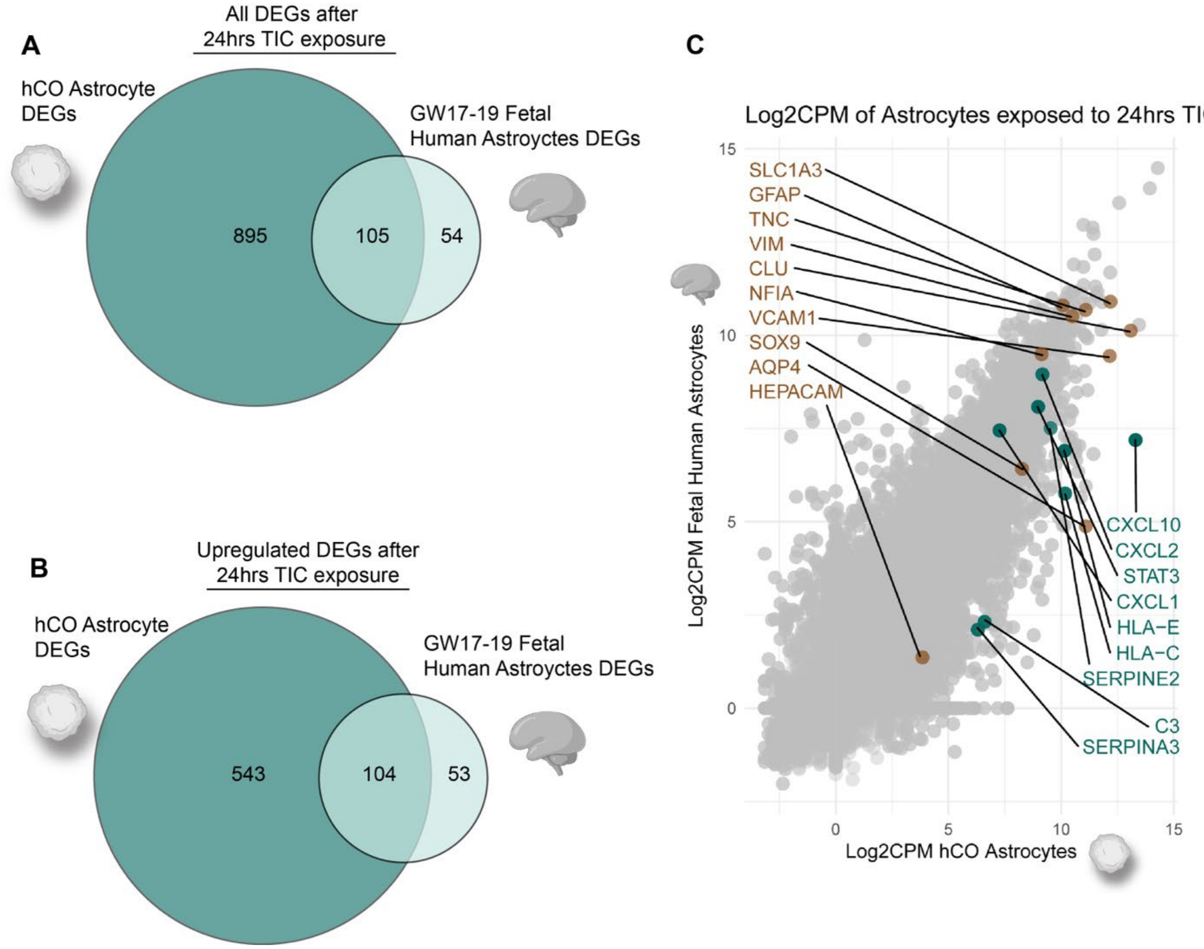


**Fig. S1. (A)** Venn diagram of all DEGs both shared and unique to astrocytes exposed to TIC for 24hrs from either hCOs or isolated from fetal tissue. **(B)** Venn diagram of only upregulated DEGs both shared and unique to astrocytes exposed to TIC for 24hrs from either hCOs or isolated from fetal tissue. **(C)** Scatterplot correlating Log2 CPM of reactive astrocytes from hCOs or fetal tissue. “Canonical” reactive astrocyte DEGs are highlighted in teal and astrocytic genes are highlighted in brown.

**Fig. S2.**

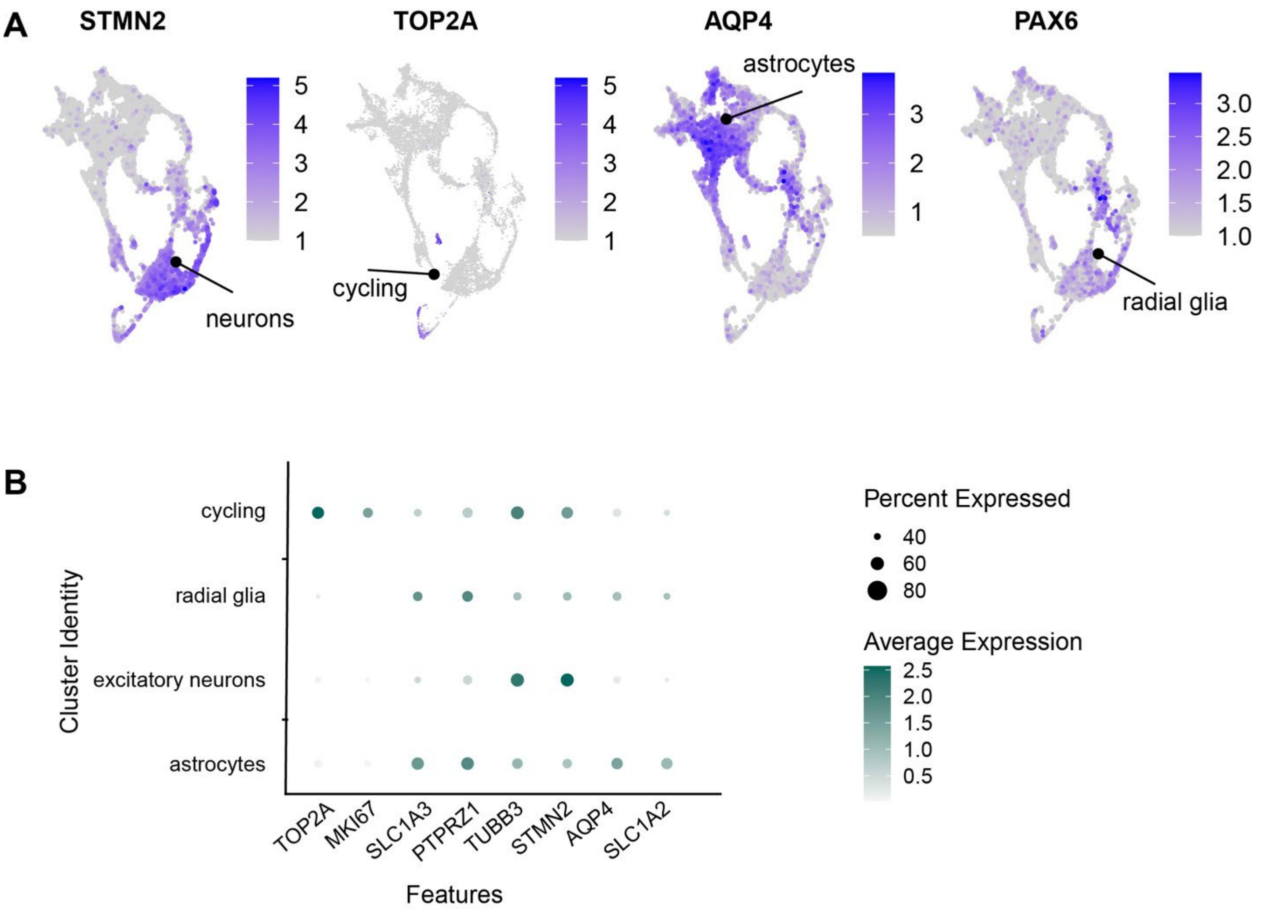


**Fig. S2. (A)** Featureplots depicting expression of cell-type specific markers for each cluster: neurons, cycling cells, astrocytes, radial glia. **(B)** Dotplot of cell-type specific marker gene expression in each named cell cluster.

**Fig. S3.**

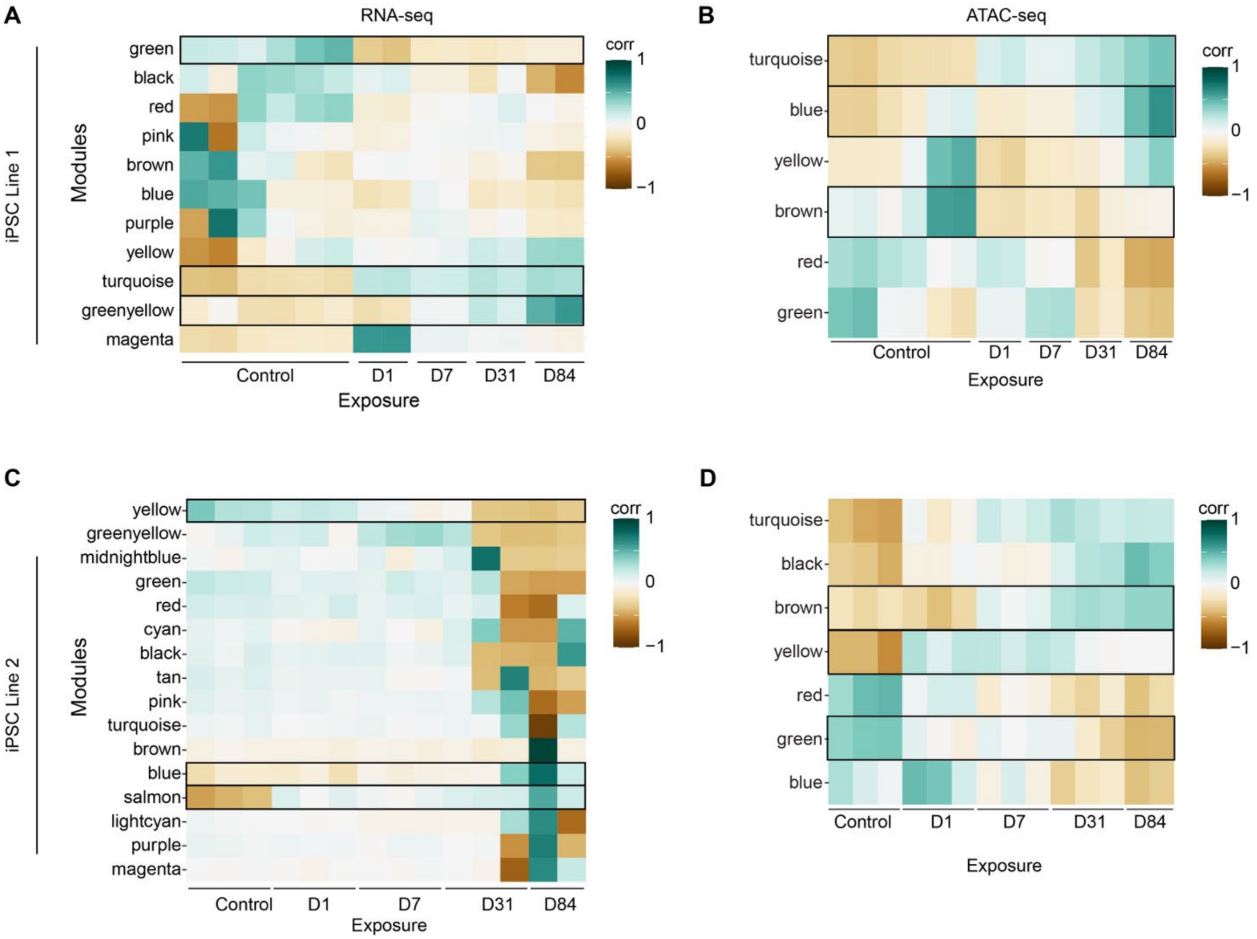


**Fig. S3. (A-D)** Complete heatmaps of WGCNA modules from both RNA-seq and ATAC-seq on purified control and reactive astrocytes at each timepoint, across 2 hiPSC lines. RNA parameters: minModuleSize=200, mergeCutHeight=0.15, ATAC parameters: minModuleSize=400, mergeCutHeight=0.15. Outlined modules refer to time-independent, time-dependent, and downregulated modules used for further analysis.

**Fig. S4.**

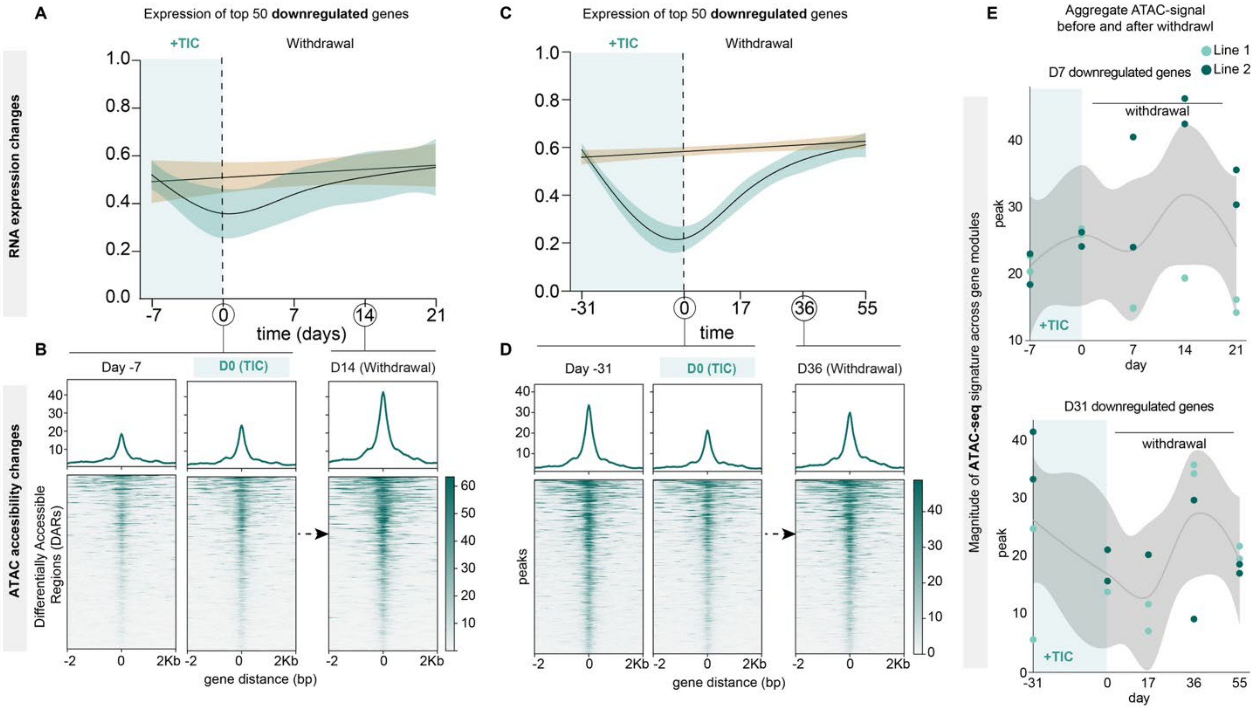


**Fig. S4. (A)** GAM plot depicting normalized Log2 CPMs of the top 50 most variably-expressed downregulated genes after acute (7 day) TIC exposure and withdrawal. **(B)** Importance plots for control, 7 day +TIC exposure, and 21 days of withdrawal from TIC exposure for time-independent gene associated DARs after acute TIC exposure and withdrawal. DARs are aligned in the middle of the plot, with +/- 2kb from the DAR. The most variably-accessible DARs are plotted toward the top. **(C)** GAM plot depicting normalized Log2 CPMs of the top 50 most variably-expressed downregulated genes after chronic (31 day) TIC exposure and withdrawal. **(D)** Importance plots for control, 31 day +TIC exposure, and 36 days of withdrawal from TIC exposure for time-dependent gene-associated DARs after acute TIC exposure and withdrawal. **(E)** Magnitude of ATAC peaks within either the time-independent or time dependent gene modules across the acute and chronic withdrawal timeline. These points correspond to the importance plots in B and D.

**Fig. S5**

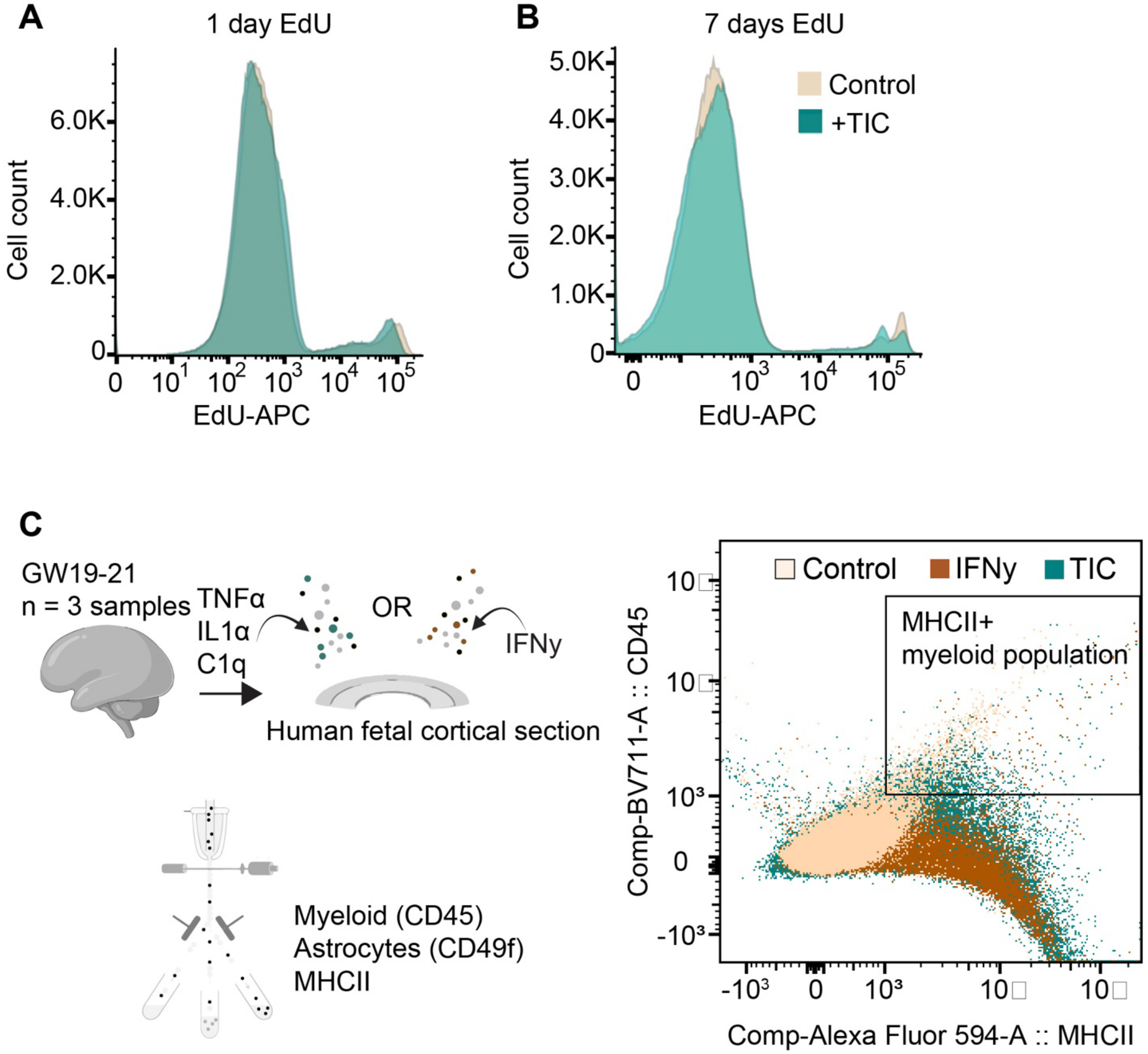


**Fig. S5. (A)** Flow analysis of hCOs cultured in normal media + EdU or media +TIC +EdU for 1 day. n=3 hCOs/condition, 12 hCOs total. **(B)** Flow analysis of hCOs cultured in normal media + EdU or media +TIC +EdU for 7 days. n=3 hCOs/condition, 12 hCOs total. **(C)** Left: 300μm GW18-21 fetal cortical sections were cultured on transwell inserts for 7 days either with no additives, +IFN-γ (2 days), or +TIC (7 days). n = 3 samples. Right: FACS plot depicting myeloid cells expressing MHCII.

**Fig. S6.**

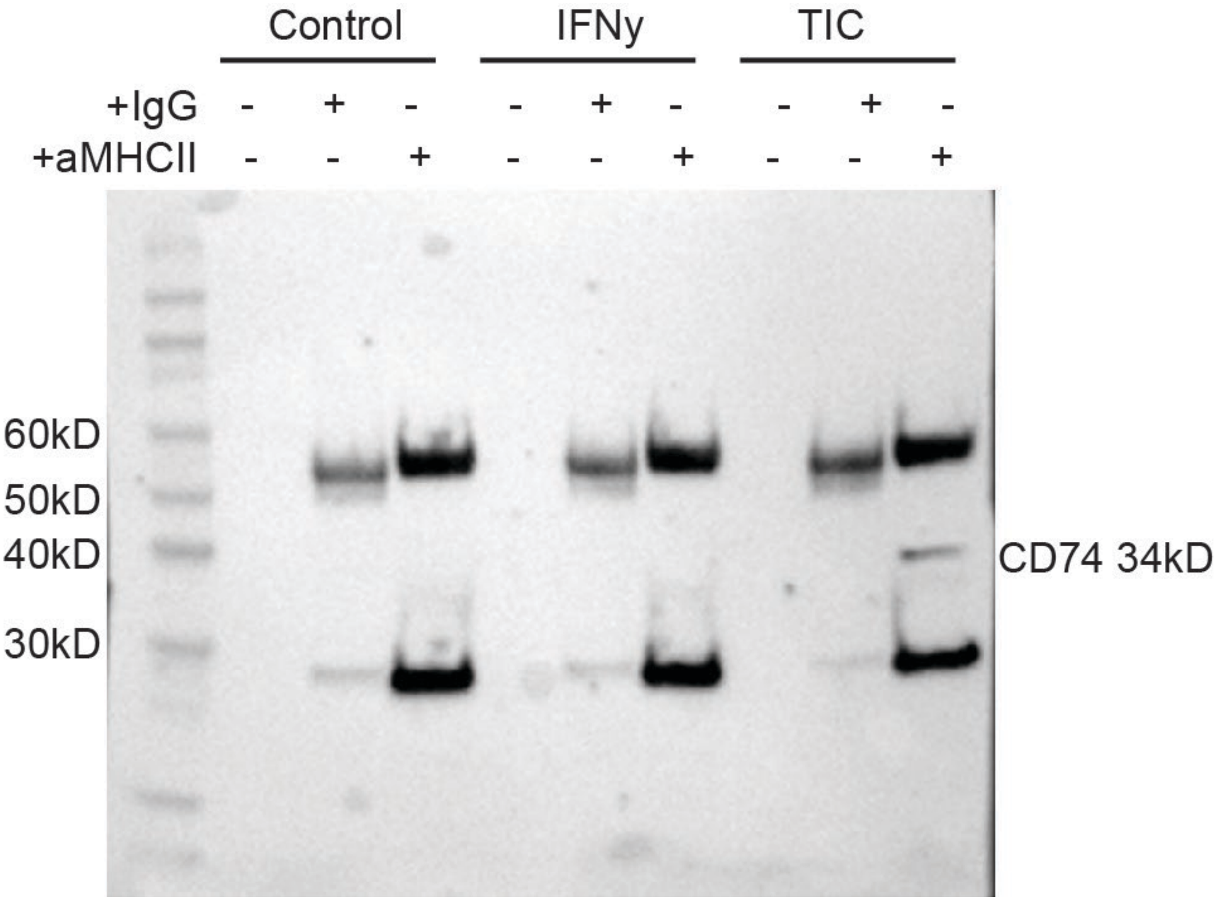


**Fig. S6.** Full Western blot of MHCII immunoprecipitation from Fig. 5J.
